# Limitations and Enhancements in Genomic Language Models: Dynamic Selection Approach

**DOI:** 10.1101/2024.11.25.624002

**Authors:** Shibo Qiu

## Abstract

Genomic Language Models (GLMs), which learn from nucleotide sequences, are crucial for understanding biological principles and excel in tasks such as sequence generation and classification. However, state-of-the-art models vary in training methods, architectures, and tokenization techniques, resulting in different strengths and weaknesses. We propose a multi-model fusion approach with a dynamic model selector that effectively integrates three models with distinct architectures. This fusion enhances predictive performance in downstream tasks, outperforming any individual model and achieving complementary advantages. Our comprehensive analysis reveals a strong correlation between model performance and motif prominence in sequences. Nevertheless, overreliance on motifs may limit the understanding of ultra-short core genes and the context of ultra-long sequences. Importantly, based on our in-depth experiments and analyses of the current three leading models, we identify unresolved issues and suggest potential future directions for the development of genomic models. The code, data, and pre-trained model are available at https://github.com/Jacob-S-Qiu/glm_dynamic_selection.

## 2 Introduction

In recent years, the rapid advancement of biotechnology has led to the successful completion of the Human Genome Project [1], resulting in an exponential growth of accessible genomic sequence data [2]. However, effectively analyzing and understanding this vast amount of genomic data to address various biomedical challenges remains a pressing issue.

Meanwhile, deep learning technologies have achieved significant breakthroughs in the field of natural language processing (NLP), including Convolutional Neural Networks (CNN) [3], Recurrent Neural Networks (RNN) [4], the Transformer architecture [5], and the BERT model [6]. These models have not only driven the progress of NLP but have also achieved remarkable success in commercial applications [7, 8].

Given the success of deep learning in NLP, researchers have begun to apply these methods to the field of biology, particularly in the analysis of nucleotide sequences. By training deep learning models on experimental data, a variety of downstream tasks have been accomplished, including:

**Genomic function prediction:** Genome-wide association studies (GWAS) predictions [9, 10], transcription factor binding site predictions [11, 12], splice site recognition [13], enhancer and silencer identification [14, 15], regulatory element identification [16], and untranslated region (UTR) predictions [17].

**Protein-related predictions:** Protein design [18], protein evolution prediction [19], protein function prediction [20, 21], and predictions of DNA/RNA-binding proteins [22].

**Gene expression and regulation:** Gene expression level predictions [23, 24], DNA methylation state predictions [25], and decoding of histone modification patterns [26].

**Structural predictions:** DNA three-dimensional structure predictions [27] and genomewide 3D folding structure predictions [28].

**Other tasks:** RNA sequencing coverage predictions [29].

Currently, genomic models are typically categorized based on training methods (e.g., Masked Language Models (MLM) [6], Conditional Language Models (CLM) [30]) or model architectures (e.g., Convolutional Neural Networks (CNN) [3], Transformer [5], and Structured State Space Models (SSM) [31]). Among these, the Transformer architecture is widely applied in genomic models [24, 32, 33, 34, 35, 36, 37, 38, 17, 39, 40, 41, 42, 43]. However, traditional position-encoded Transformers face performance bottlenecks when handling long sequences, typically supporting sequence lengths of around 1 kb. To address this limitation, Rotary Position Embeddings [44] have been introduced, extending sequence lengths to approximately 10 kb and demonstrating excellent performance in downstream tasks [45, 46]. Furthermore, genomic language models based on the BERT architecture [47, 48, 49, 40], as well as CNN- and RNN-based models [50, 51], have also been developed. Recently, SSM-based models for long-sequence modeling have extended the sequence length capacity to over 100 kb [52, 53, 54, 55, 56], offering new possibilities for processing ultra-long genomic sequences.

In machine learning and pattern recognition, dynamic selection (DS) methods have been proposed to integrate the strengths of various algorithms[57]. These methods have achieved remarkable performance in multiple classifier systems (MCS) [58, 59, 60, 61]. Dynamic selection dynamically selects the most suitable classifier for a given input based on its features by leveraging a diverse set of base classifiers. The effectiveness of MCS relies on the diversity of the base classifiers; if the classifiers provide overly similar features, the combined result may be suboptimal [62].

Inspired by dynamic selection, this study proposes a multi-model architecture based on dynamic selection to enhance downstream task performance. We selected three models with significant architectural differences as base classifiers to achieve feature extraction diversity. These models are Hyena [53], NTv2 [45], and CD-GPT [35], whose core architectures are SSM, Rotary Position Encoding-based Transformer, and Position Encoding-based Transformer, respectively. Their maximum sequence processing capacities are 160 kb, 12 kb, and 1 kb. Each model achieves excellent performance in its specialized downstream tasks, with some attaining state-of-the-art (SoTA) levels [53, 45, 35].

By constructing a base classifier pool consisting of these three models, we effectively leverage their complementary strengths in feature extraction. Furthermore, we improved these three models so that they can not only serve as base classifiers but also function as dynamic selectors, dynamically selecting the most suitable model among the three base classifiers to execute specific tasks. Experiments were conducted on downstream tasks with sequence lengths of 500 bp and 20,000 bp, demonstrating that the dynamic selector model outperformed any single model.

Additionally, as our model architecture is based on dynamic selection from multiple advanced models, we conducted an in-depth analysis of the error causes for each dynamic selector and base classifier model in different downstream tasks. While most genomic model researchers tend to focus on achieving higher benchmark scores, detailed analyses of sequence processing capabilities across different genomic models are often neglected. Through sequence-level and overall analysis of each downstream task, we identified several issues with current genomic models and proposed directions for future optimization.

### Contributions of This Study

- **Proposed a dynamic multi-model architecture** that integrates the capabilities of diverse base classification models with varying architectures. The dynamic selection mechanism enables the framework to outperform individual base classifiers, achieving state-of-the-art (SoTA) performance in downstream genomic tasks.
- **Conducted a comprehensive comparative analysis** of results from multiple models, identifying key differences in their performance. Additionally, performed an indepth error analysis of sequences prone to misclassification for each model, revealing architectural limitations in existing genomic model designs.
- **Identified pivotal, unresolved questions for genomic models**, grounded in our comprehensive experiments and critical reflections on current model constraints. These insights extend beyond the immediate scope of our findings, offering valuable guidance for future research. By highlighting specific avenues of exploration and potential pitfalls, we aim to help researchers focus their efforts more effectively, thus laying the foundation for deeper investigations into genomic sequence modeling.

## 3 Results

### 3.1 Performance on Short Sequence Tasks

#### 3.1.1 Model Prediction Performance

To evaluate the performance of each model on short sequence tasks, we conducted experiments on DNA sequences with a length of 500 bp. This length was chosen to test the classification capability of the models at single-nucleotide resolution. We utilized the Human Enhancers Cohn module from the Genomic Benchmarks dataset [63], which contains experimentally validated human enhancer sequences. In the experiments, we compared the performance of the three base classifier models (Hyena, NTv2, and CD-GPT) and their corresponding dynamic selector models. Table 1 summarizes the accuracy and F1-scores for all models.

**Figure 1:**
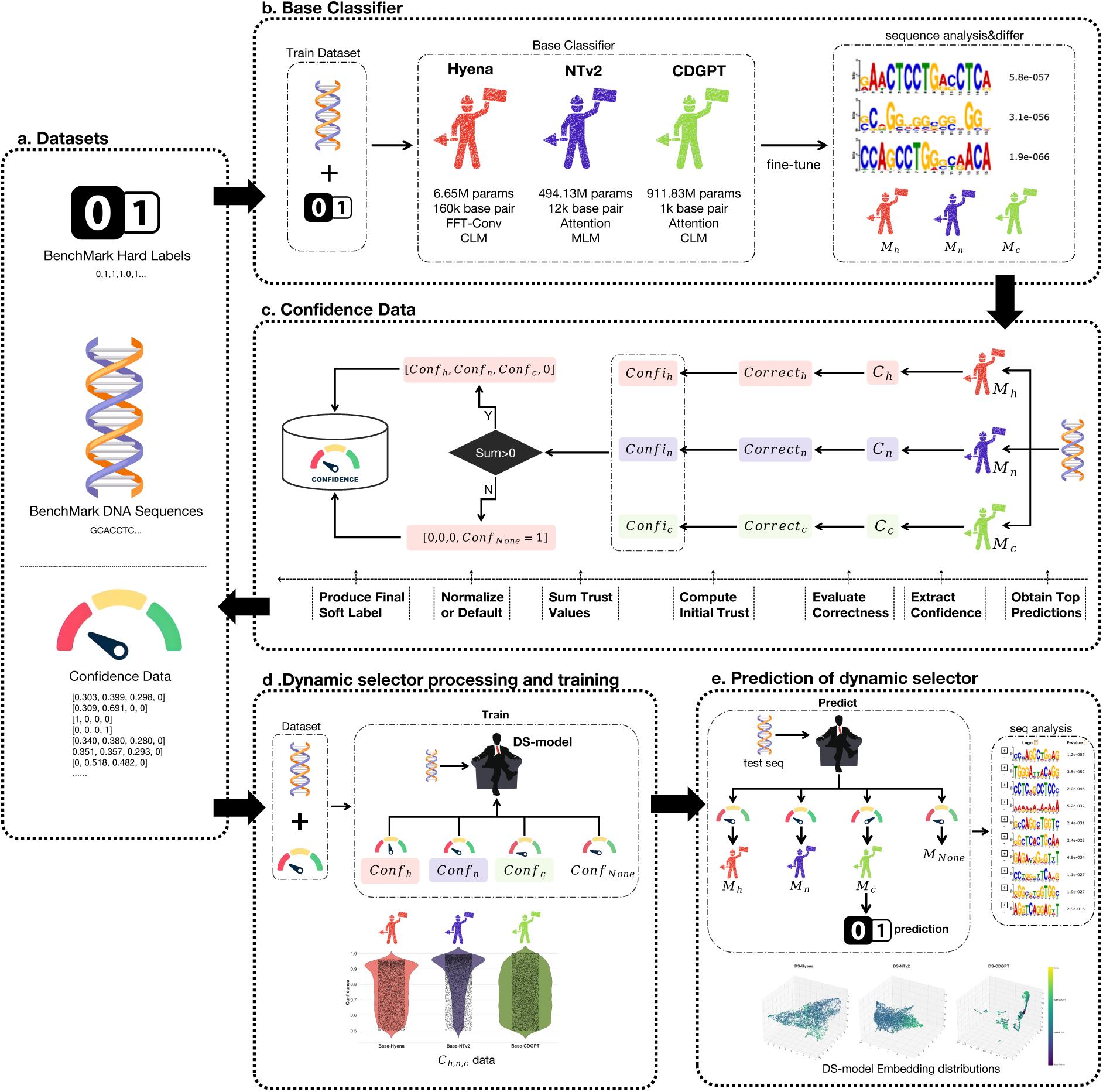
Overall workflow of the proposed method. (**a**) The dataset used throughout the experiment includes *Hard Labels* and *Sequences* obtained from the benchmark, while *Confidence* values are derived from subsequent experiments. (**b**) We fine-tune three base models and perform differential sequence analysis on their outputs. (**c**) The *Confidence Dataset* is constructed by extracting each model’s maximum output and aggregating the results through a series of processing steps. (**d**) We train the dynamic selector using both the input sequences and the confidence scores from the three base classifiers. (**e**) The dynamic selector’s predictions are evaluated, and sequence alignment is conducted on the final outputs to further assess its performance.

**Table 1:**
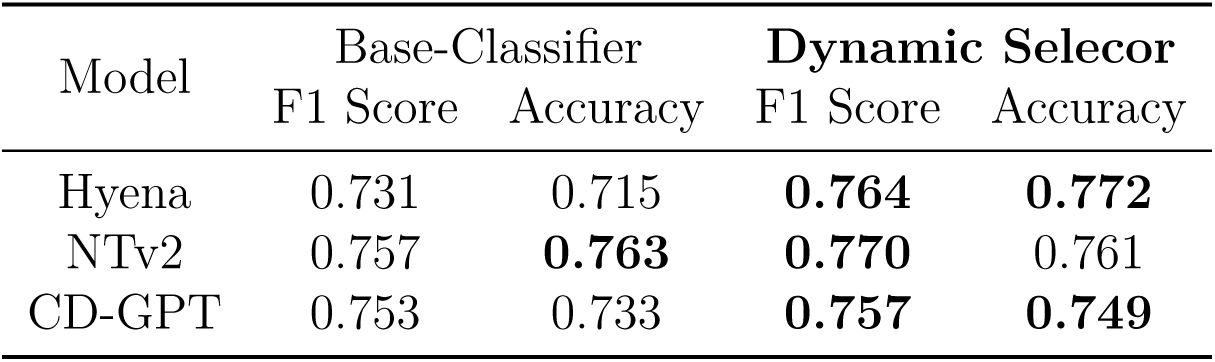
Performance comparison between base classifier models and dynamic selector models on 500 bp short sequence tasks.

| Model | Base-Classifier |  | Dynamic Selector |  |
| --- | --- | --- | --- | --- |
|  | F1 Score | Accuracy | F1 Score | Accuracy |
| Hyena | 0.731 | 0.715 | **0.764** | **0.772** |
| NTv2 | 0.757 | **0.763** | **0.770** | 0.761 |
| CD-GPT | 0.753 | 0.733 | **0.757** | **0.749** |

As shown in the table, the dynamic selector models outperform their corresponding base classifier models in both accuracy and F1-score. This indicates that the dynamic selector can effectively integrate the strengths of different models, enhancing predictive performance in short sequence tasks.

#### 3.1.2 Task Allocation Analysis of Dynamic Selectors

To further understand the working mechanism of the dynamic selector models, we analyzed the task allocation statistics for each dynamic selector across the base classifiers. The results are presented in Table 2.

**Table 2:** Task allocation statistics for dynamic selector models on 500 bp short sequence tasks.

| Selector | Classifier | Total Count | TP+TN | FP+FN | Accuracy |
| --- | --- | --- | --- | --- | --- |
| DS-Hyena | Base-Hyena | 1 | 1 | 0 | 100% |
|  | Base-NTv2 | 5325 | 4083 | 1242 | 76.7% |
|  | Base-CDGPT | 1622 | 1222 | 400 | 75.3% |
|  | None | 0 | 0 | 0 | — |
| DS-NTv2 | Base-Hyena | 0 | 0 | 0 | — |
|  | Base-NTv2 | 5830 | 4364 | 1466 | 74.9% |
|  | Base-CDGPT | 1118 | 919 | 199 | 82.2% |
|  | None | 0 | 0 | 0 | — |
| DS-CDGPT | Base-Hyena | 220 | 204 | 16 | 92.7% |
|  | Base-NTv2 | 2254 | 1703 | 551 | 75.6% |
|  | Base-CDGPT | 4420 | 3287 | 1133 | 74.4% |
|  | None | 54 | 41 | 13 | 75.9% |

From the table, it can be observed that the dynamic selector models in short sequence tasks tend to allocate most of the tasks to NTv2 and CD-GPT, both of which are attention-based models. Specifically, NTv2 and CD-GPT models handled approximately 98% of the tasks on average, while the Hyena model was responsible for only about 2%. This aligns with the superior performance of NTv2 and CD-GPT in processing short sequences, demonstrating that the dynamic selector can reasonably assign tasks to the most suitable base classifier based on sequence characteristics.

#### 3.1.3 Visualization Analysis of Dynamic Selector Models

To further validate the above conclusions, we performed dimensionality reduction on the embedding vectors of the dynamic selector models, and the results are shown in Figure 2.

**Figure 2:**
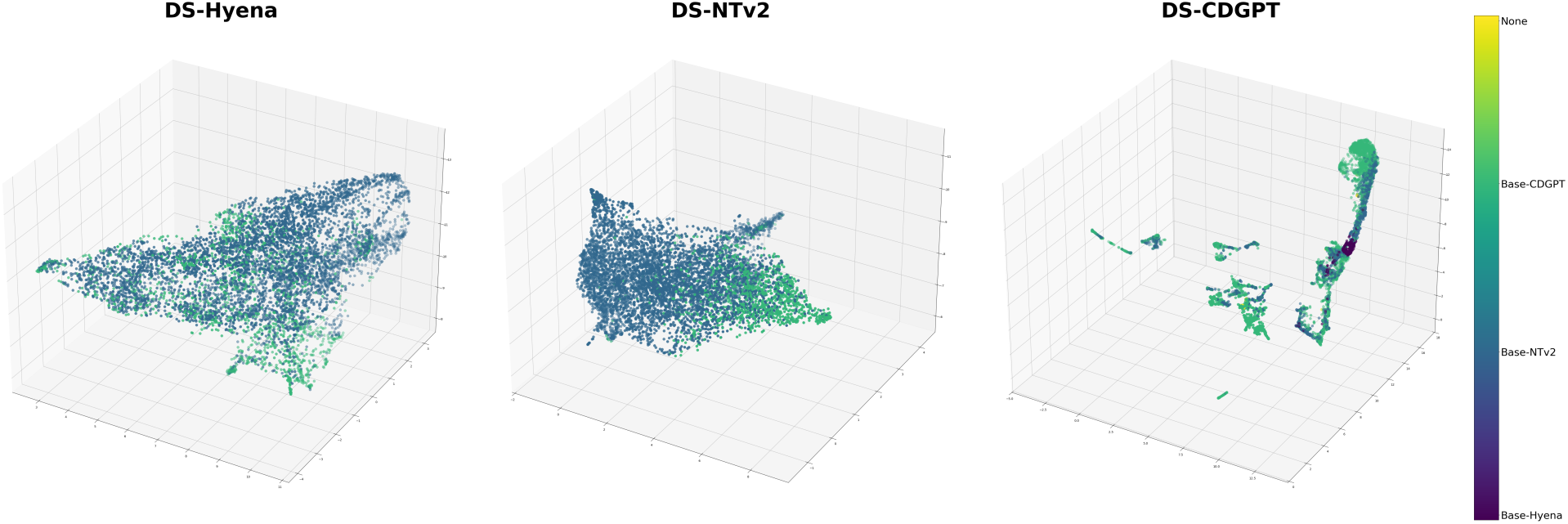
Embedding distributions of dynamic selector models on 500 bp short sequence tasks.

From the figure, it can be observed that the sequences form distinct clusters in the embedding space, and the dynamic selector model assigns them to different base classifiers based on sequence characteristics. Most of the sequences in DS-Hyena and DS-NTv2 models are assigned to NTv2 and CD-GPT, further supporting the results from the task allocation statistics.

#### 3.1.4 Correlation Between Sequence Features and Model Performance

To explore the relationship between model predictive performance and sequence features, we conducted motif analysis on the prediction results of the DS-NTv2 model. Using the MEME tool [64], we compared the true positive (TP), false positive (FP), true negative (TN), and false negative (FN) sequences predicted by the model.

The sequence analysis results for NTv2 are shown in Figure 7. It can be seen that TN and FN sequences exhibit high similarity in key motifs, and these motifs have significantly low E-values, indicating high enrichment in the sequences. This enrichment may mislead the model’s feature judgment to some extent.

**Figure 7:**
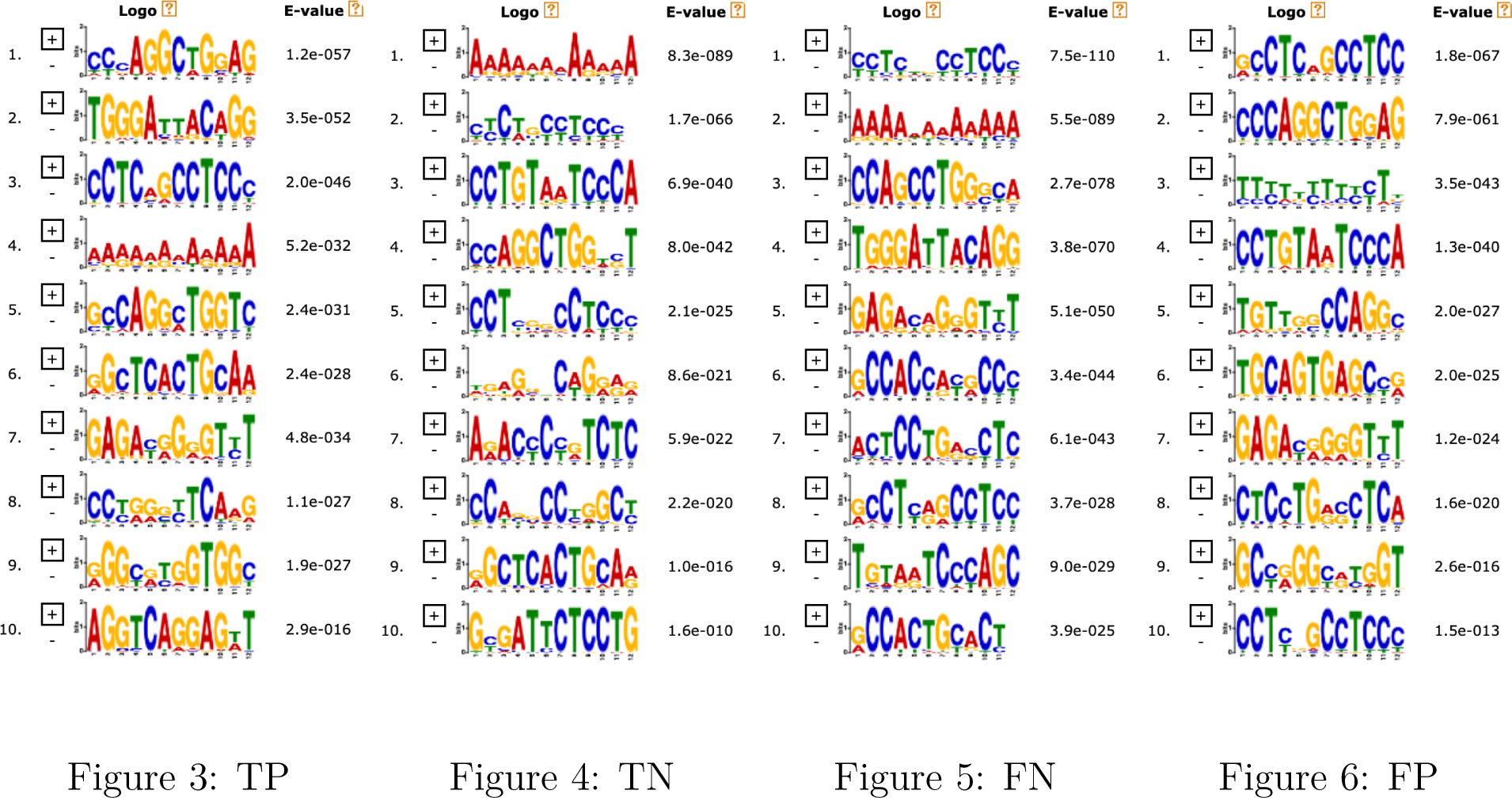
Motif analysis results of the Base-NTv2 classifier under the DS-NTv2 model

The motif analysis results for the CD-GPT classifier are shown in Figure 12. TP and FP sequences exhibit similar motif characteristics, indicating that the model relies heavily on these significant motifs when predicting positive sequences. This reliance also leads to misclassifications for FP sequences. Thus, our findings are reinforced: the model’s misclassification is highly correlated with the significance (E-value) of motifs in the sequences.

**Figure 12:**
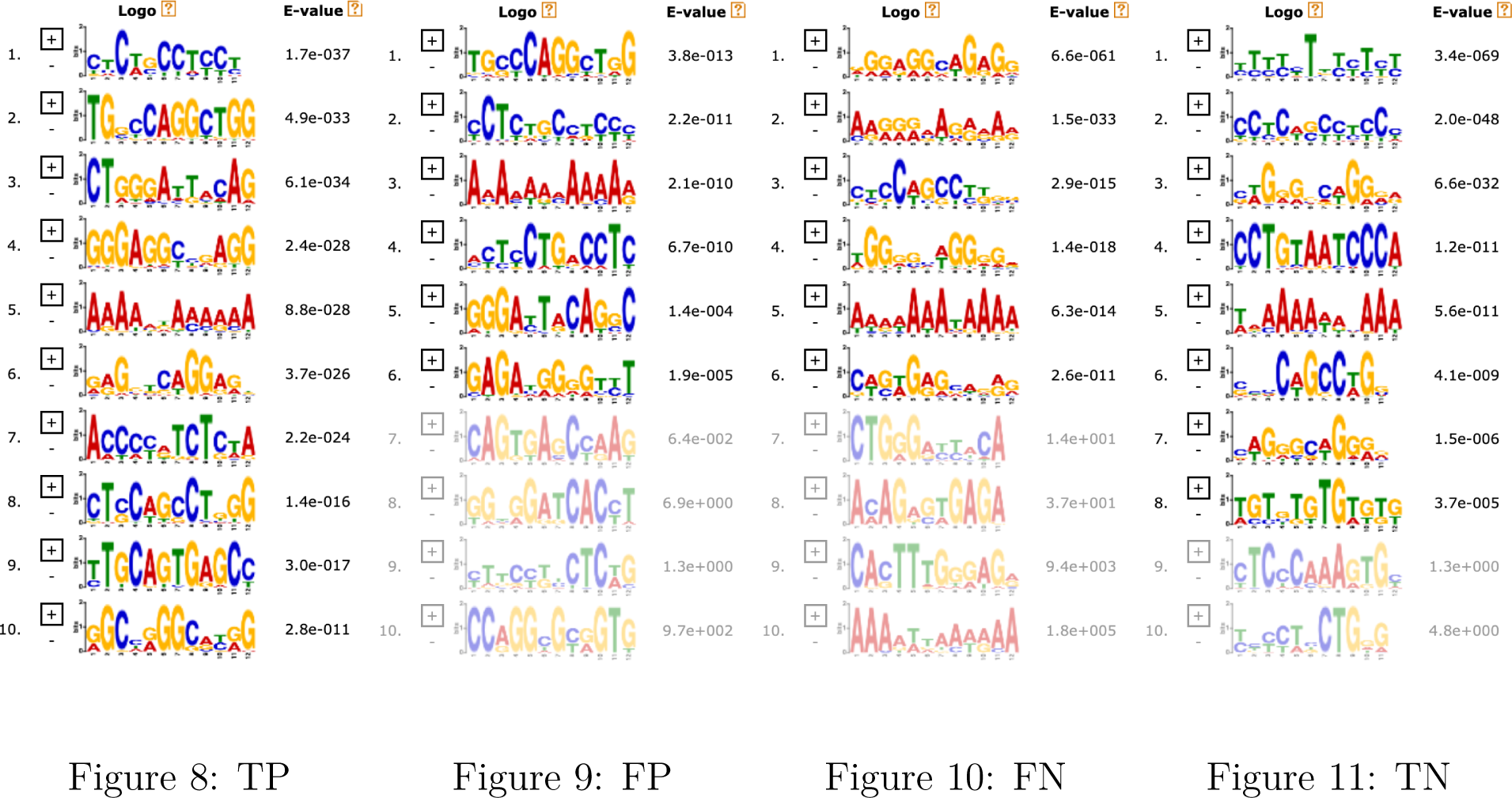
Motif analysis results of the Base-CDGPT classifier under the DS-NTv2 model

##### Summary

The above figures illustrate the motif distribution across different prediction categories. It is evident that similar motifs play a significant role in model misclassifications. Specifically, when negative sequences contain motifs similar to those in positive sequences, the model is more likely to produce misclassifications. This highlights the strong correlation between the model’s predictive performance and the significance of motifs in the sequences.

### 3.2 Performance on Long Sequence Tasks

#### 3.2.1 Model Prediction Performance

To evaluate the performance of each model on long sequence tasks, we conducted experiments on DNA sequences with a length of 20,000 bp. This sequence length was chosen to test the models’ ability to handle long-range dependencies and global sequence features, simulating real gene expression regulatory environments. We utilized gene expression data from the GM12878 cell line [65], with sequence ranges covering 10,000 bp upstream and downstream of transcription start sites (TSS).

Due to the input sequence length limitations of each model, sequences were appropriately truncated (details provided in Appendix Table 9).

Table 3 summarizes the performance of the base classifier models and their corresponding dynamic selector models on long sequence tasks. It is evident that dynamic selector models outperform their respective base classifiers in both accuracy and F1-score, indicating that dynamic selectors effectively integrate the strengths of each model to enhance performance in long sequence tasks.

**Table 3:** Performance comparison between base classifier models and dynamic selector models on 20,000 bp long sequence tasks.

| Model | Base-Classifier |  | Dynamic Selector |  |
| --- | --- | --- | --- | --- |
|  | F1 Score | Accuracy | F1 Score | Accuracy |
| Hyena | 0.820 | 0.818 | **0.839** | **0.837** |
| NTv2 | 0.835 | 0.836 | **0.841** | **0.839** |
| CD-GPT | 0.803 | 0.783 | **0.842** | **0.844** |

As shown in the table, although the CD-GPT model is constrained by its input length when handling long sequences, its overall performance is significantly improved with the assistance of the dynamic selector. This demonstrates that dynamic selectors effectively allocate tasks in long sequence tasks, fully leveraging the strengths of each base classifier.

#### 3.2.2 Task Allocation Analysis of Dynamic Selector Models

To further understand the working mechanism of dynamic selector models in long sequence tasks, we analyzed the task allocation statistics for each dynamic selector across the base classifiers. The results are presented in Table 4.

**Table 4:** Task allocation statistics for dynamic selector models on 20,000 bp long sequence tasks.

| Selector | Classifier | Total Count | TP+TN | FP+FN | Accuracy |
| --- | --- | --- | --- | --- | --- |
| DS-Hyena | Base-Hyena | 179 | 151 | 28 | 84.4% |
|  | Base-NTv2 | 809 | 676 | 133 | 83.6% |
|  | Base-CDGPT | 2 | 2 | 0 | 100% |
|  | None | 0 | 0 | 0 | — |
| DS-NTv2 | Base-Hyena | 201 | 185 | 16 | 92.0% |
|  | Base-NTv2 | 784 | 638 | 146 | 81.4% |
|  | Base-CDGPT | 5 | 5 | 0 | 100% |
|  | None | 0 | 0 | 0 | — |
| DS-CDGPT | Base-Hyena | 176 | 153 | 23 | 86.9% |
|  | Base-NTv2 | 762 | 632 | 130 | 82.9% |
|  | Base-CDGPT | 52 | 49 | 3 | 94.2% |
|  | None | 0 | 0 | 0 | — |

From the table, it is evident that in long sequence tasks, the dynamic selector models predominantly allocate tasks to Base-Hyena and Base-NTv2, both of which are capable of handling longer sequences. Specifically, Base-Hyena and Base-NTv2 models jointly handled approximately 93% of the tasks, while Base-CDGPT was responsible for only about 7%. This aligns with the superior performance of Base-Hyena and Base-NTv2 in processing long sequences, demonstrating that the dynamic selector can reasonably assign tasks based on sequence length and features. Additionally, it is notable that while Base-CDGPT handled a significant portion of tasks in short sequence processing, its role diminished significantly in long sequence tasks.

#### 3.2.3 Visualization Analysis of Dynamic Selector Models

To further validate the above conclusions, we performed dimensionality reduction on the embedding vectors of the dynamic selector models. The results are shown in Figure 13.

**Figure 13:**
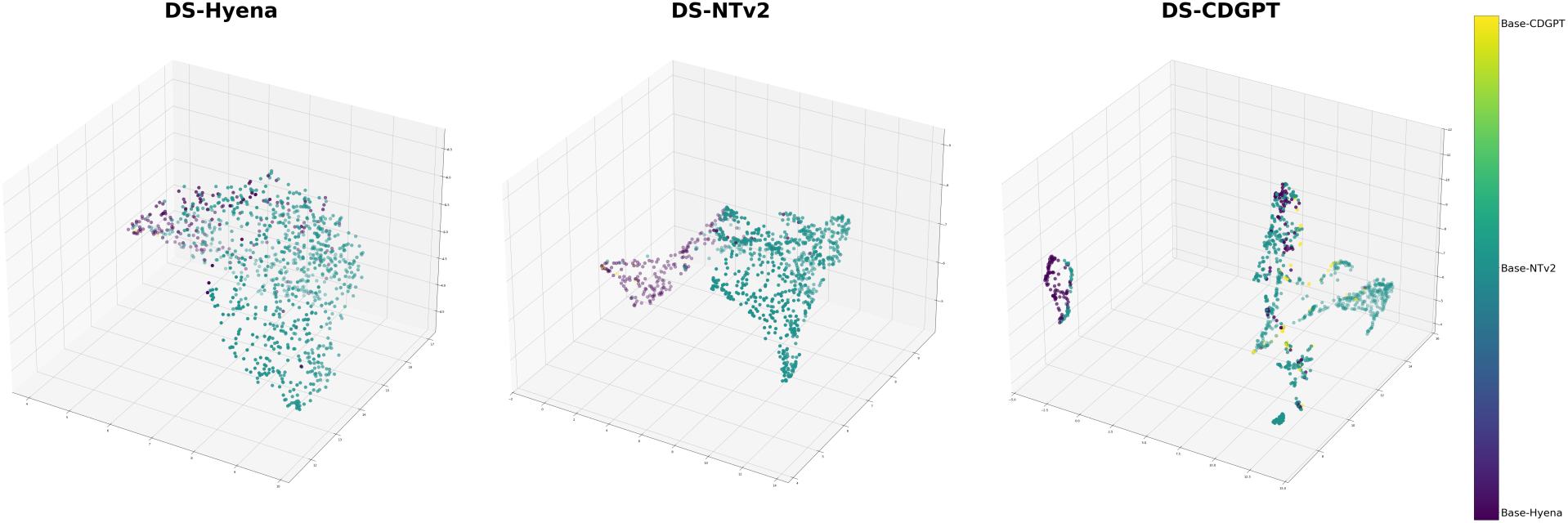
Embedding distributions of dynamic selector models on 20,000 bp long sequence tasks.

From the figure, it can be observed that sequences form distinct clusters in the embedding space, and the dynamic selector models assign them to different base classifiers based on sequence features. Most sequences in DS-Hyena and DS-NTv2 models are allocated to Base-Hyena and Base-NTv2 models, further supporting the results from the task allocation statistics.

#### 3.2.4 Detailed Analysis of Model Prediction Results

To gain deeper insights into model behavior in long sequence tasks, we performed a detailed analysis of the prediction results for each base classifier model. Based on prediction correctness, sequences in the test set were categorized into the following four groups:

1. **All base classifiers predict correctly:** All three models correctly predicted the sequence.
2. **At least two base classifiers predict correctly:** Two models predicted correctly, and one model predicted incorrectly.
3. **Only one base classifier predicts correctly:** Only one model predicted correctly, while the other two models predicted incorrectly.
4. **All base classifiers predict incorrectly:** All three models predicted the sequence incorrectly.

Table 5 presents the number and proportion of sequences in each category.

**Table 5:**
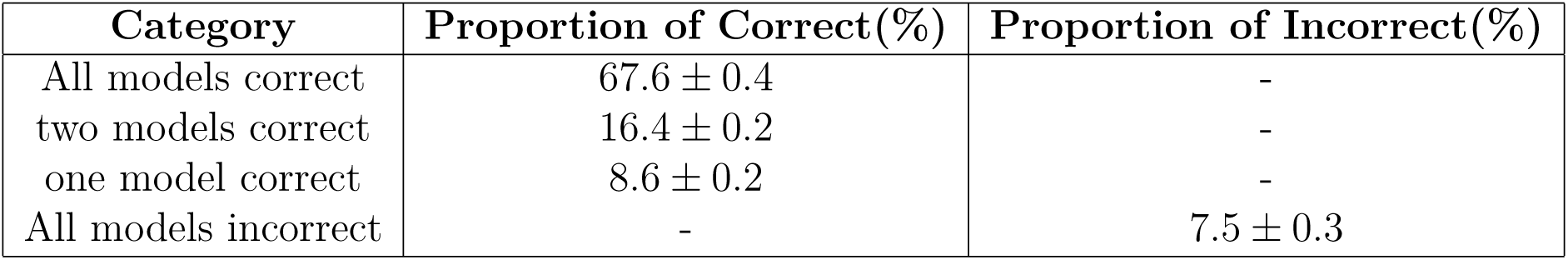
Categorical statistics of prediction results on the test set.

| Category | Proportion of Correct(%) | Proportion of Incorrect(%) |
| --- | --- | --- |
| All models correct | $67.6 \pm 0.4$ | - |
| two models correct | $16.4 \pm 0.2$ | - |
| one model correct | $8.6 \pm 0.2$ | - |
| All models incorrect | - | $7.5 \pm 0.3$ |

#### 3.2.5 Correlation Between Sequence Features and Model Performance

To investigate the impact of sequence features on model predictive performance, we used the MEME Suite to perform motif analysis on sequences from different prediction correctness categories. Specifically, motif discovery and significance analysis were conducted for the four categories, and the results are shown in Figure 14.

**Figure 14:**
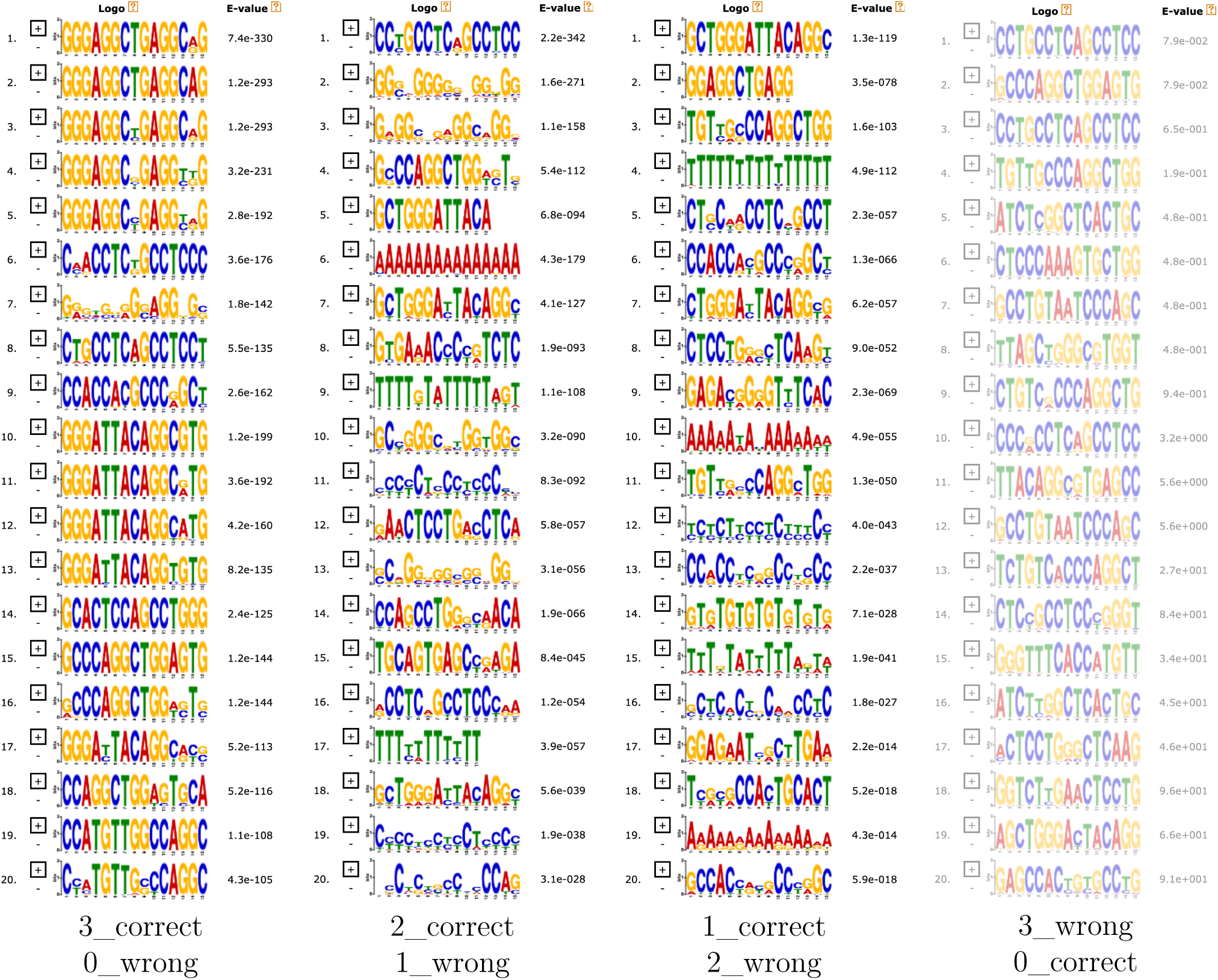
Motif significance analysis of sequences with different prediction correctness categories.

The analysis results indicate:

- **Sequences where all models predict correctly:** These sequences contain highly significant motifs with extremely low E-values, indicating strong biological features. The models effectively capture these features, leading to correct predictions.
- **Sequences where some models predict correctly:** As the number of correct predictions decreases, the motif significance in the sequences also diminishes. The E-values increase, suggesting that the frequency or conservation of motifs in these sequences is lower, making it more challenging for the models to identify these features.
- **Sequences where all models predict incorrectly:** These sequences exhibit the lowest motif significance, and statistically significant motifs may even be undetectable. For this scenario, we propose two hypotheses:

1. **Hypothesis 1:** The diminished motif significance in these sequences reduces the models’ ability to make accurate predictions.
2. **Hypothesis 2:** There may be unique factors in these sequences where the true determinants of expression are not related to motif significance. Instead, they might lie on **other clues**, such as longer-range context or in a few ultra-short core genes. However, the models’ **inability to identify** these critical features leads to decreased prediction accuracy.

### 3.3 In-depth Analysis of Error-Prone Sequences

To further investigate the relationship between model prediction accuracy and motif significance in long sequence tasks, we referred to [66], which highlighted that data in the hg38 coordinate system contains more feature information compared to the hg19 coordinate system. Thus, we upgraded the DNA sequence data in the current dataset from the hg19 coordinate system to the hg38 coordinate system for the same gene IDs (gene_id) and repeated the experiments.

First, we selected all sequences previously classified as 3_wrong sequences, where all base classifier models predicted incorrectly (see Figure 14), and obtained their corresponding sequences in the hg38 coordinate system for motif analysis. The results, shown in Figure 16, revealed that the motif significance of these sequences in the hg38 coordinate system was greatly enhanced, with E-values decreasing from the 10^−2^ magnitude to the 10^−469^ magnitude, indicating much stronger motif characteristics.

**Figure 15:**
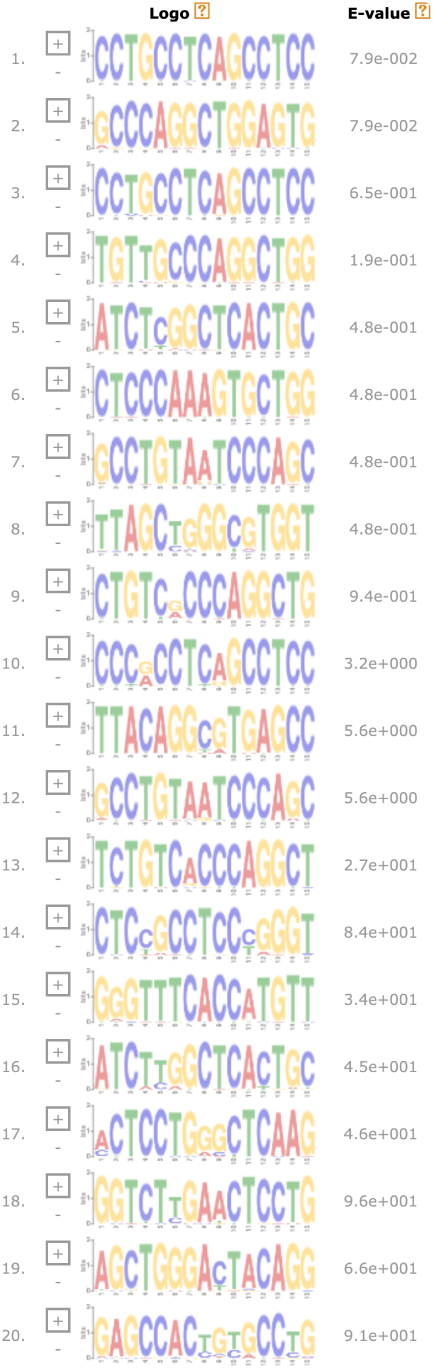
Same as the rightmost in Fig 14(for better comparison).

**Figure 16:**
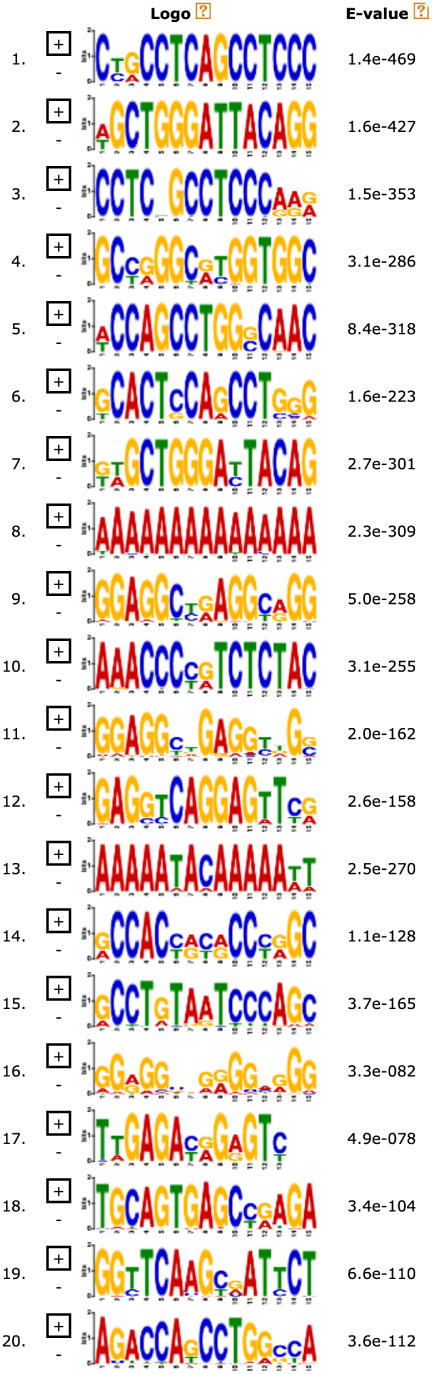
Sequences with the same gene_id in hg38 and redetect feature.

Based on our earlier **Hypothesis 1**, enhanced motif significance should improve model prediction capabilities. However, the experimental results (see Table 6) indicate that despite the significant increase in motif significance for error-prone sequences under the hg38 system, the overall model accuracy did not improve and even slightly declined.

**Table 6:** Performance of base classifiers and selector models on hg38 data.

| Model | Base-Classifier |  | Dynamic Selector |  |
| --- | --- | --- | --- | --- |
|  | F1 Score | Accuracy | F1 Score | Accuracy |
| Hyena | 0.749 | 0.711 | **0.781** | **0.752** |
| NT-V2 | 0.770 | **0.756** | **0.780** | 0.751 |
| CD-GPT | 0.777 | **0.764** | **0.782** | 0.754 |

To investigate the underlying reasons, we reanalyzed the sequences that were misclassified by all base classifiers in the hg38 coordinate system. Figure 17 shows the motif characteristics of these sequences, revealing a strong similarity to the enhanced error-prone sequences in the hg19 coordinate system (Figure 16). This suggests that regardless of the coordinate system, these sequences pose similar challenges for the current models, even when their features are enhanced.

**Figure 17:**
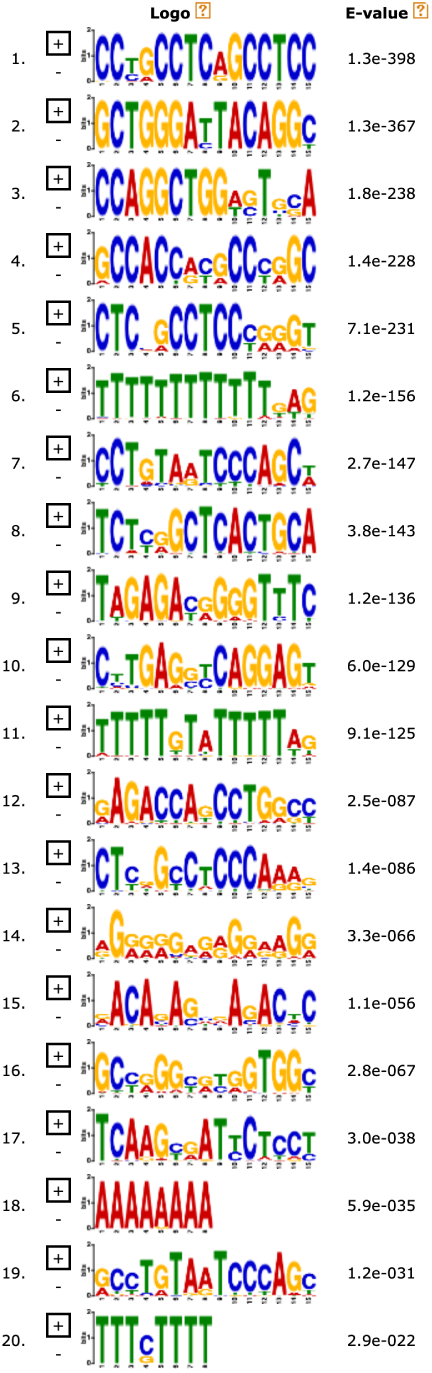
Error-prone sequences feature after training on hg38 data.

These results suggest that even when models can recognize more significant motif features, their predictive performance for certain sequences does not necessarily improve and may even deteriorate. A potential cause of this performance decline is that the enhanced importance of error-prone motifs in hg38 might lead the model to misclassify sequences it previously predicted correctly in hg19. This implies that **Hypothesis 1** alone cannot fully explain these misclassifications. Instead, **Hypothesis 2** may be the key factor contributing to the model’s errors, suggesting that the expression regulation of these sequences might not rely on significant motif features and **the models may be suspected of overbelieving motif features**. Furthermore, **these models are unable to effectively identify this critical information, and the length of this information does not fall within typical motif regions**, such as:

1. **Long-distance contextual information (>10 kbp):** The expression of the sequence might be influenced by remote regulatory elements beyond the model’s receptive field.
2. **Ultra-short core sequences (1–6 bp):** Critical regulatory elements might consist of extremely short sequences that, despite their crucial importance, the model struggles to capture.
3. …

## 4 Discussion

This study proposes a dynamic selector-based multi-model fusion architecture that leverages three leading and structurally distinct models—Hyena, NTv2, and CD-GPT—as base classifiers. Dynamic selectors allocate tasks among these classifiers to enhance downstream performance. Experimental results demonstrate that the **dynamic selector achieves better performance than any single base classifier on downstream tasks(SoTA)**, which holds significant value for specific application scenarios. However, this architecture requires task-specific fine-tuning, leading to high resource consumption. Therefore, when efficiency and cost-effectiveness are priorities, this approach may not be ideal.

An in-depth analysis of the capabilities of the base classifiers and dynamic selector models revealed a strong correlation between model performance and the significance of motifs in sequences. Given that the three selected models represent state-of-the-art architectures, this finding underscores significant progress in the ability of current genomic models to recognize certain biological sequence features. However, these models still exhibit critical limitations: **they may give a excessive trust for motif’s feature and more likely to misunderstand sequence when the key information falls outside typical motif lengths**.

Even the EVO model, which leads in overall performance, has deficiencies in generating coherent ultra-long sequences and identifying ultra-short critical genes [56]. Another innovative architecture, ChatNT, achieved high performance (more than 0.95) in 6 out of 18 tasks but underperformed (less than 0.7) in 8 tasks [67]. These observations align closely with our Hypothesis 2. Therefore, we advocate for some researchers to dedicate more effort toward long-sequence research rather than short-sequence model, as it holds greater value. The reasoning is straightforward: if a model is limited to processing sequences of only 1,000 base pairs (bp), then no matter how intelligent or adept it is at understanding and recognizing biological significance, it will remain oblivious to the information beyond that limit— the 1,001st base pair and onward. Conversely, by extending our efforts to sequences of 100 kbp, we enable models to learn and integrate the full spectrum of information embedded within these longer sequences. It becomes merely a matter of time or some special technical structures before such models achieve a comprehensive understanding.

**Therefore, many deep learning models in genome sequence analysis can indeed automatically learn highly characteristic motifs (e.g., transcription factor binding sites, splice sites) from training data, and to some extent, these motifs correspond to experimentally validated functional regions. However, they merely extract features (even with attention architectures, the effect is limited); more importantly, enhancing the model’s understanding capabilities should be the focus. Pure sequence models cannot encompass protein interactions, spatiotemporal expression, evolutionary information, and other factors that ultimately determine biological functions and phenotypes. Having only sequence or limited structural information prevents the model from forming a systemic biological understanding; it is necessary to incorporate literature knowledge and database annotations (such as gene functions, protein domain information, protein interaction networks, etc.) into the model’s training or reasoning processes.**

To develop more effective biological models, **it is essential to explore more appropriate and accessible training data, design comprehensive model training frameworks, and prioritize the research of genome models capable of effiiciently extracting features from ultra-long sequences.** By addressing these three critical challenges, we can achieve significantly improved biological model performance.

### 4.1 Outstanding Question

1. How can we establish a unified, high-quality training benchmark dataset that directs researchers toward more effective model training and evaluation? Is the current field of genomic modeling lacking a practical and impactful dataset comparable to ImageNet?
2. How can we effectively integrate multi-omics data (e.g., transcriptomics, epigenomics, proteomics) to enhance the model’s understanding of biological processes in their entirety? Achieving a comprehensive view of life activities likely depends on the ability to synthesize insights across multiple biological data modalities.
3. How should we define and measure a genomic model’s “understanding”of biological meaning? Which technical approaches could facilitate the transition from mere feature extraction to deeper biological semantic comprehension? Would creating specialized “intermediate Module”for feature interpretation and output be necessary?
4. In genomic research, how can we design a universal architecture suitable for multi-species, multi-omics data, thereby addressing the heterogeneity and diversity inherent in genomic datasets? Should the model’s capacity for effective transfer learning be considered an essential criterion for evaluation?

## Appendix: Methods

### A.1 Base Classifier Processing

#### A.1.1 Selection of Base Classifier Models

To construct a base classifier pool capable of adapting to diverse DNA sequence characteristics, this study selected three high-performing genomic large models: CD-GPT, NT-V2, and Hyena. These models each excel in handling sequences of varying lengths and capturing sequence features, complementing one another to improve overall predictive performance. The specific reasons for choosing these three models are detailed below.

The CD-GPT model is based on the traditional Transformer architecture [35], specializing in processing short DNA sequences (e.g., fragments up to 1 kb). As a Conditional Language Modeling (CLM) model, it is suitable for generation and classification tasks. Using Byte Pair Encoding (BPE) tokenization, it effectively captures local features in short sequences, such as regions around Transcription Start Sites (TSS).

The NT-V2 model employs a Masked Language Modeling (MLM) architecture [45], focusing on capturing features in medium-length genomic sequences. The MLM architecture trains the model through the prediction of randomly masked fragments and utilizes nonoverlapping k-mer tokenization, enabling efficient processing of genomic sequences. Additionally, it employs Rotary Embeddings, which mitigate the decay of positional information in long sequences, allowing it to capture dependencies in sequences up to 12 kb.

Finally, HyenaDNA is an innovative long-sequence language model [53] designed for ultralong DNA sequences (20 kbp and beyond, up to 160 kbp). Using a hybrid architecture combining autoregressive CLM and sequence-to-sequence modeling (SSM), it excels in both generation and feature extraction for ultra-long sequences. Furthermore, its novel use of Fast Fourier Transform (FFT)-based long convolution mechanisms replaces traditional attention mechanisms, reducing computational complexity from *O*(*n*^2^) to *O*(*n* log *n*). This makes it well-suited for identifying remote regulatory regions (e.g., enhancer-TSS associations) and parsing sequences at the whole-genome level.

#### A.1.2 Accuracy Testing of Base Classifier Models

During the initial testing phase of the base classifier model set, the three models were trained and fine-tuned on different datasets to evaluate their capabilities in gene expression prediction for specific task scenarios. Let the accuracy of model *M_i_* be *A_i_*, where *i* ∈ {*c, n, h*} represents the CD-GPT, NT-V2, and Hyena models, respectively.

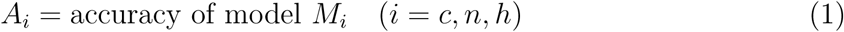

To further quantify the prediction confidence of the models, we extracted the maximum output value from the classification head of each base classifier model, referred to as the confidence score *C_i_*. This is defined as:

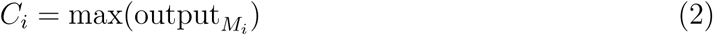

Here, output*_M_* represents the prediction output of model *M_i_* for a given input sequence. The confidence score *C_i_* reflects the confidence of model *M_i_* in processing the current sequence and serves as a key input for constructing the task selector model. By analyzing the predictive capabilities of each model, we developed a highly generalized multi-model task processing architecture. This enables the task selector model to assign each sequence to the most suitable classifier model based on sequence characteristics, thereby improving overall prediction performance.

### A.2 Confidence Set Extraction

To effectively train the task selector model, this study constructed a confidence set (Meta Dataset). The confidence set includes the prediction confidence scores of each base classifier model for every sequence, representing the models’confidence in their predictions. Specifically, the construction process involves extracting confidence scores from the classifier models’ outputs and generating soft labels based on these scores. These soft labels reflect the relative strengths of each classifier model in handling sequences within its domain of expertise.

#### A.2.1 Confidence Extraction and Calculation

For a given DNA sequence x*_i_*, we input it into the three base classifier models: *M_c_* (CD-GPT), *M_n_* (NT-V2), and *M_h_* (Hyena). For each model, the output output*_M_* (x*_i_*) is obtained, where *j* ∈ {*c, n, h*} denotes the respective models.

The output output*_M_* (x*_i_*) is a vector containing the model’s confidence scores for different classes. The maximum value of this vector is defined as the prediction confidence *C_j_*(x*_i_*) of model *M_j_* for sequence x*_i_*:

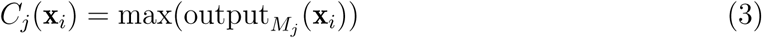

The confidence score *C_j_*(x*_i_*) measures the confidence of model *M_j_* in its prediction for sequence x*_i_*. A higher value indicates greater confidence in the prediction result.

During the construction of the confidence set, this study extracted the confidence values *C_c_*(x*_i_*), *C_n_*(x*_i_*), and *C_h_*(x*_i_*) for each classifier model and each data point, creating a foundational dataset containing the prediction confidences of all models.

### A.3 Construction of Confidence Labels

When generating confidence labels, one-hot encoding was not used; instead, soft labels were employed to enhance the generalization capability of the model. Specifically, for each sequence x*_i_*, if the sequence was correctly predicted by model *M_j_*, its confidence score was retained for constructing the soft label. If the model’s prediction was incorrect, the corresponding confidence score was set to zero. Additionally, for cases where all classifier models predicted incorrectly, a value of 1 was assigned to Confi(None) to represent the “all wrong” scenario.

The soft label vector **Conf**(x*_i_*) is defined as:

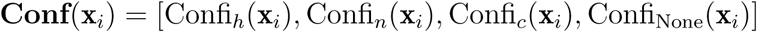

The calculation method is as follows:

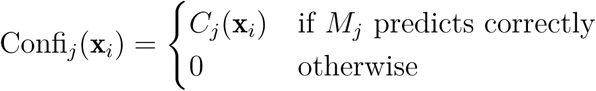

For cases where *M_h_*, *M_n_*, and *M_c_* all predict incorrectly:

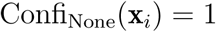

To maintain the probabilistic distribution characteristics of the soft labels, the labels were normalized. The final confidence label for each sequence **Conf**(x*_i_*) is normalized using the following formula:

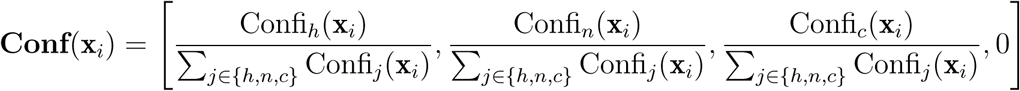

For scenarios where all classifier models predict incorrectly, we set:

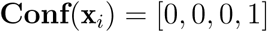

The soft label dataset constructed using the above method preserves the relative confidence distributions of each base classifier model in their respective areas of expertise. This dataset better reflects the matching between sequence features and model capabilities, which is critical for training the task selector model.

### A.4 Training of the Dynamic Selector Model

This section describes the training process for the Dynamic Selector Model. The goal of the selector model is to allocate sequences to the most suitable base classifier model based on their features, thereby improving overall prediction accuracy. To achieve optimal task allocation, we designed a specialized loss function to ensure that the output probability distribution of the selector model aligns as closely as possible with the true soft label probability distribution.

#### A.4.1 Loss Function Design

The training objective of the dynamic selector model is to make its output task allocation probability distribution 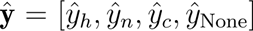 match the true soft label probability distribution y = [*y_h_, y_n_, y_c_, y*_None_] as closely as possible. Here, 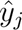 represents the confidence assigned by the selector model to the *j*-th classifier model, and *y_j_* represents the corresponding confidence in the true soft label, where *j* ∈ {*h, n, c,* None} denotes the Hyena model, NT-V2 model, CD-GPT model, and the “all wrong” case, respectively.

To achieve this, we employ the **Kullback-Leibler Divergence (KLD)** as the loss function. KLD measures the relative entropy between two probability distributions, quantifying the difference between the selector model’s predicted probability distribution 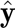 and the true probability distribution y. The KLD is defined as:

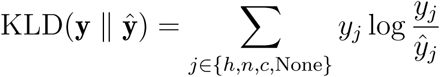

In this equation:

- *y_j_* is the true soft label probability for the *j*-th category.
- 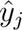 is the predicted probability by the selector model for the *j*-th category.
- The KLD encourages the selector model to minimize the divergence between its predictions and the true soft labels, guiding it to make accurate task assignments.

#### A.4.2 Training Procedure

During training, each sequence x*_i_* is passed through the selector model to obtain the predicted task allocation probabilities 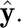 The KLD between the predicted distribution 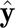 and the true soft label distribution y is computed and used as the loss for optimization. The training process iteratively updates the selector model’s parameters to minimize the KLD loss, ensuring that the model learns to allocate tasks effectively based on sequence characteristics.

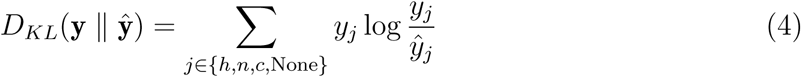

During training, we minimize the sum of KL divergences over all training samples to ensure that the output of the dynamic selector model progressively aligns with the soft label distribution. Assuming the training set contains *N* DNA sequences, with each sequence having a predicted distribution 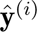 and a true soft label y^(^*^i^*^)^, the overall loss function *L* is defined as:

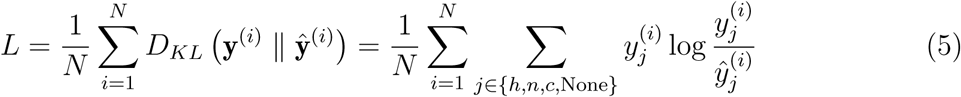

Here, 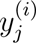 and 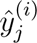 represent the confidence of the *j*-th classifier model for the *i*-th sequence in the true distribution and predicted distribution, respectively. A key feature of KL divergence is its asymmetry, i.e., 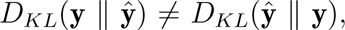 which makes it more sensitive to high-confidence categories in the true distribution y (i.e., larger *y_j_* values). This property directs the selector model to focus on improving the accuracy of predictions for high-confidence categories.

#### A.4.3 Training Optimization of the Dynamic Selector Model

The dynamic selector model minimizes the KL divergence loss function, driving its output probability distribution to closely match the true soft label distribution. During training, the Adam optimizer is employed to update the parameters of the selector model and minimize the loss function *L*. The optimized selector model can then make optimal task allocation decisions across the classifier models based on the characteristics of the input sequence. The overall logic of the method is outlined in Algorithm 1.

## B Appendix: Supplementary Materials

### B.1 Datasets and Pretrained Models

#### Training Datasets

The dataset for short-sequence experiments was obtained from the Genomic Benchmarks dataset [63], specifically the *human_enhancers_cohn* subset. Detailed information about this dataset is provided in Table 9. The long-sequence experiments used gene expression data from the human GM12878 cell line under the hg19 coordinate system [65].

#### Pretrained Models

To ensure reproducibility, the pretrained models used in this study are accessible via the following links:

- Hyena: https://huggingface.co/LongSafari/hyenadna-medium-160k-seqlen-hf
- NTv2: https://huggingface.co/InstaDeepAI/nucleotide-transformer-v2-500m-multi-speci
- CD-GPT: https://github.com/TencentAI4S/CD-GPT

### B.2 Training Parameters

#### B.2.1 Training Parameters for Base Classifier Models

Table 7 lists the hyperparameter configurations for the three base classifier models. Each model adopts tailored hyperparameters during training to achieve optimal task adaptation performance.

**Table 7:**
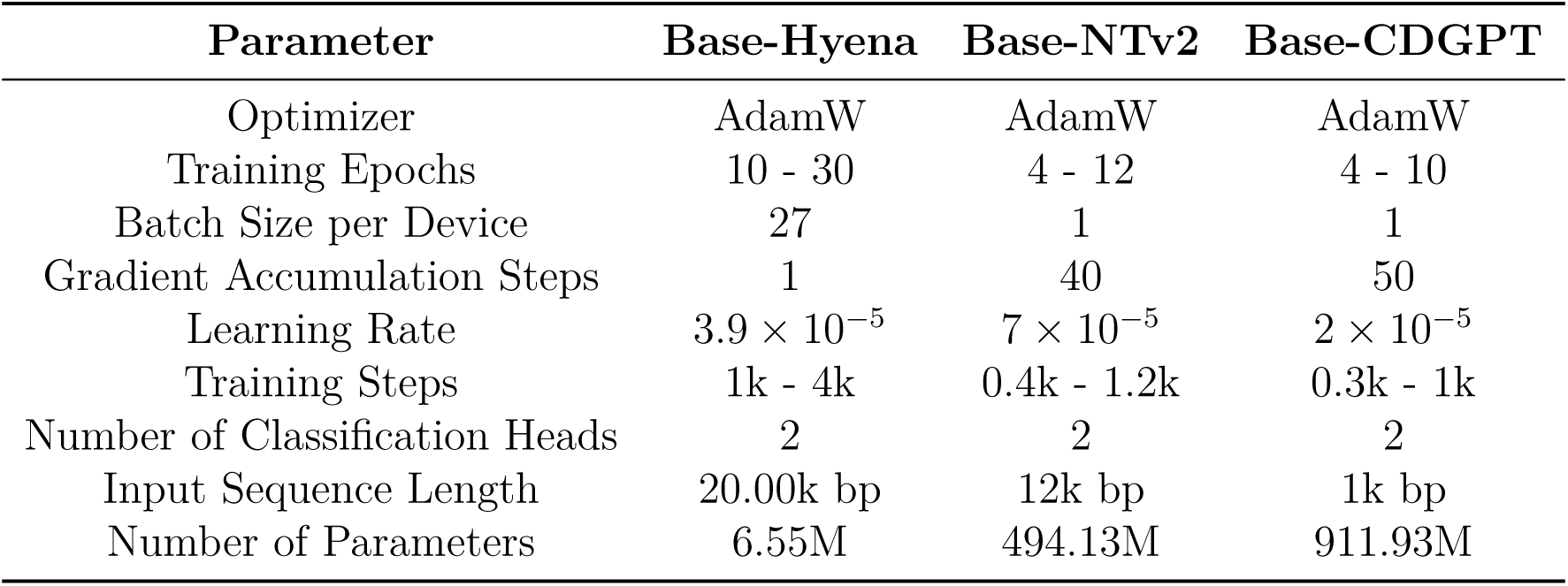
Hyperparameter Settings for Base Classifier Models.

##### Algorithm 1

Dynamic Selector Model Training

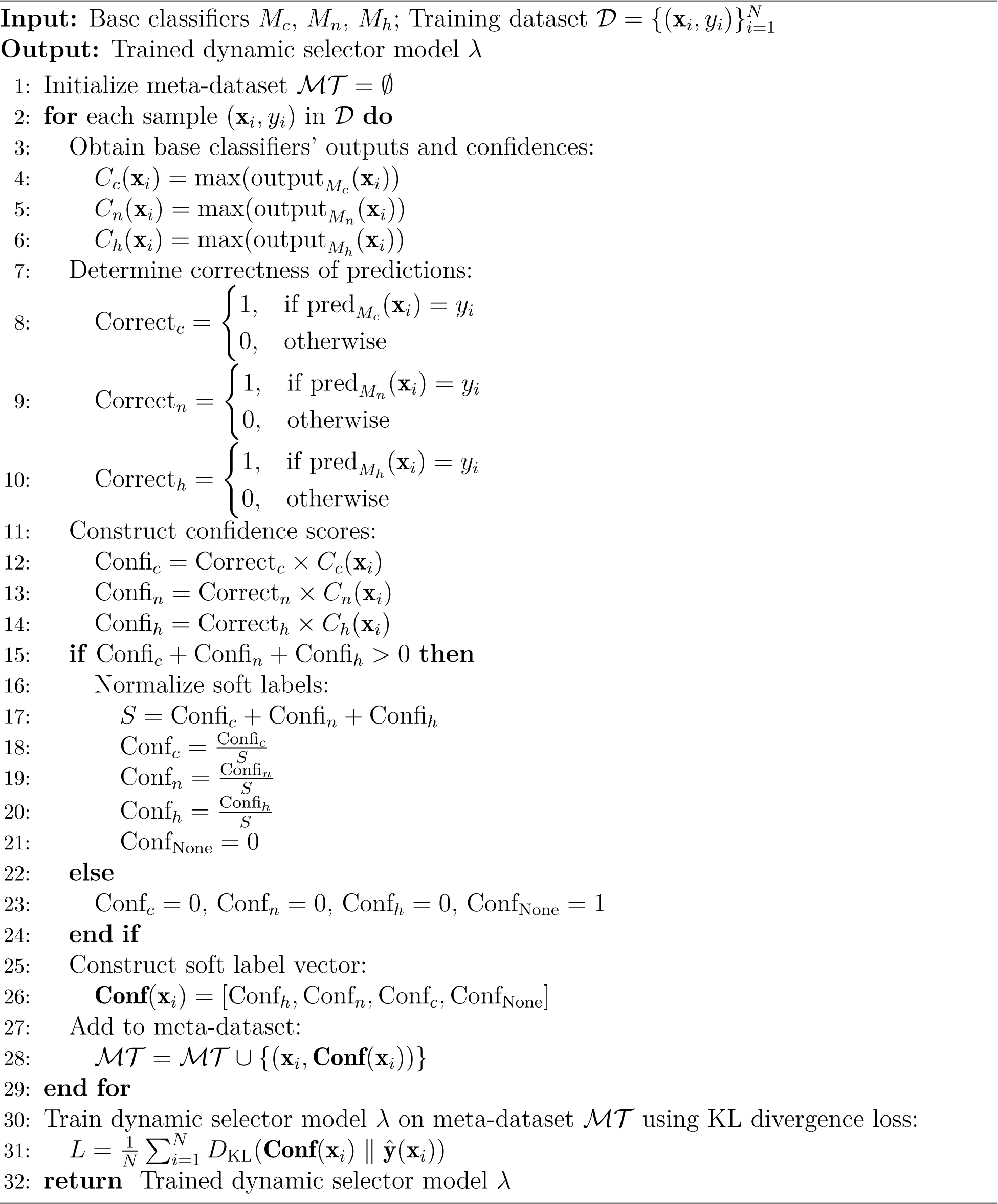

#### B.2.2 Training Parameters for Dynamic Selector Models

Table 8 outlines the hyperparameter configurations for the dynamic selector models. To ensure effective learning of task allocation strategies, fine-tuned parameter adjustments were performed for each model.

**Table 8:**
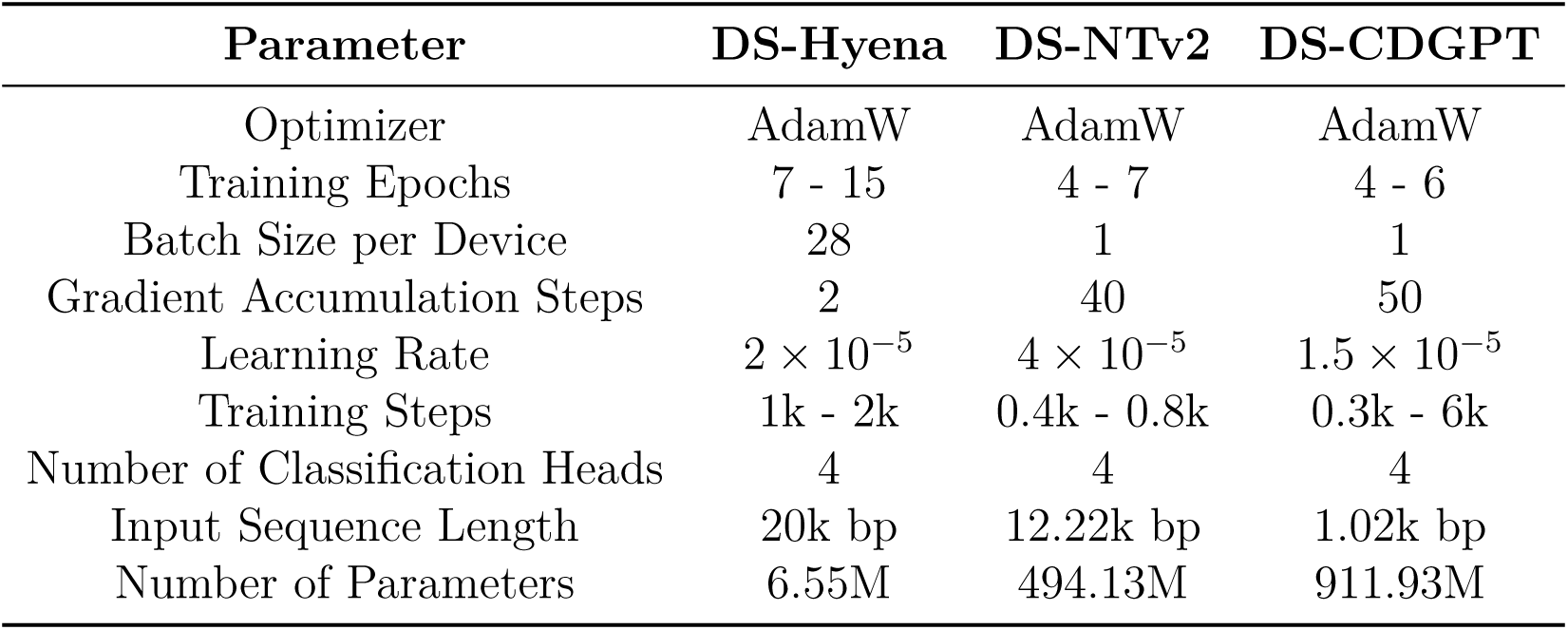
Hyperparameter Settings for Dynamic Selector Models.

| Parameter | DS-Hyena | DS-NTv2 | DS-CDGPT |
| --- | --- | --- | --- |
| Optimizer | AdamW | AdamW | AdamW |
| Training Epochs | 7 - 15 | 4 - 7 | 4 - 6 |
| Batch Size per Device | 28 | 1 | 1 |
| Gradient Accumulation Steps | 2 | 40 | 50 |
| Learning Rate | $2 \times 10^{-5}$ | $4 \times 10^{-5}$ | $1.5 \times 10^{-5}$ |
| Training Steps | 1k - 2k | 0.4k - 0.8k | 0.3k - 6k |
| Number of Classification Heads | 4 | 4 | 4 |
| Input Sequence Length | 20k bp | 12.22k bp | 1.02k bp |
| Number of Parameters | 6.55M | 494.13M | 911.93M |

### B.3 MEME Sequence Analysis Parameters

For the motif analysis, we employed the MEME Suite tool and set different parameters for short and long sequences to account for variations in sequence length and characteristics. The detailed parameter settings are as follows:

- Short Sequence Tasks:

~~~
meme {fastafile} -dna -oc . -nostatus -time 14400 -mod anr -nmotifs 20
-minw 6 -maxw 15 -objfun classic -revcomp -markov_order 0 -p 10
~~~
- Long Sequence Tasks:

~~~
meme {fastafile} -dna -oc . -nostatus -time 14400 -mod zoops -nmotifs 10
-minw 8 -maxw 12 -objfun classic -revcomp -markov_order 0 -p 10
~~~

### B.4 Supplementary Figures and Tables

The following supplementary figures and tables provide additional support for the analysis and discussion of the main results presented in this study.

#### B.4.1 Model Performance Comparison Table

Table 9 presents a comparison of the three models in terms of parameter count, maximum acceptable sequence length, and input lengths for short and long sequence tasks.

**Table 9:** Model Performance Comparison Table.

| Metric | Hyena | NTv2 | CDGPTA |
| --- | --- | --- | --- |
| Maximum Input Sequence Length | 160,000 bp | 12,288 bp | 1,024 bp |
| Number of Hidden Neurons | 256 | 1,024 | 2,304 |
| Parameter Count | 6.65 M | 494.13 M | 911.83 M |
| Input Length for Short Tasks | 500 bp | 500 bp | 500 bp |
| Input Length for Long Tasks | 20,000 bp | 12,288 bp | 1,024 bp |
| Position Relative to TSS | [-10,000,10,000] | [-10,000,2,288] | [-974,50] |

**Table 10:** Prediction Result Statistics of Base Classifier Models on 500bp Short Sequence Tasks. We conducted pairwise comparisons between models to identify potential patterns. In the Combined Accuracy row, the probability of both Model1 and Model2 making correct predictions is the highest (69.3%), which potentially indicates that the architectures of Model1 and Model2 are relatively similar. Furthermore, it suggests that these two models are more adept at handling short-sequence tasks.

|  | Base-CDGPT(1) | Base-NTv2(2) | Base-Hyena(3) |
| --- | --- | --- | --- |
| **Input Sequence Length** | 0.5k | 0.5k | 0.5k |
| **Individual Accuracy** | 73.1% ± 0.1% | 75.4% ± 0.3% | 71.2% ± 0.3% |
| **Combined Performance** | All models being correct: 61.9% ± 0.3% | | |
| | All models being wrong: 14.5% ± 0.4% | | |
| **Combined Accuracy** | Accuracy of Models 1 and 2: 69.1% ± 0.2% | | |
| | Accuracy of Models 1 and 3: 65.3% ± 0.2% | | |
| | Accuracy of Models 2 and 3: 65.0% ± 0.3% | | |
| **Combined Error Rate** | Error Rate of Models 1 and 2: 17.7% ± 0.2% | | |
| | Error Rate of Models 1 and 3: 18.6% ± 0.1% | | |
| | Error Rate of Models 2 and 3: 17.3% ± 0.4% | | |

#### B.4.2 Prediction Statistics of Base Classifier Models

#### B.4.3 Supplementary Figures

The following supplementary figures were generated during the experiments, including the distribution of loss values, confidence scores, and visualizations of the embedding space.

**Table 11:** Prediction Result Statistics of Base Classifier Models on 20kbp Long Sequence Tasks. In the Combined Accuracy row, the number of tasks correctly predicted by both Model1 and Model3 is approximately 69.4%, which is the lowest among all pairwise comparisons. This indicates that the architectural differences between the two models are more pronounced in handling long-sequence tasks. Additionally, in the Combined Error Rate row, the probability of both models making incorrect predictions is also the lowest at only 9.4%. These observations suggest that the differences between these two models, such as architectural design or input sequence length handling, result in relatively distinct task processing capabilities. It is precisely these subtle differences that the dynamic selector is designed to exploit.

|  | Base-CDGPT(1) | Base-NTv2(2) | Base-Hyena(3) |
| --- | --- | --- | --- |
| **Input Sequence Length** | 1k | 12k | 20k |
| **Individual Accuracy** | $82.0\% \pm 0.2\%$ | $83.5\% \pm 0.2\%$ | $80.2\% \pm 0.3\%$ |
| **Combined Performance** | All models being correct: $67.6\% \pm 0.4\%$<br>All models being wrong: $7.5\% \pm 0.3\%$ | | |
| **Combined Accuracy** | Accuracy of Models 1 and 2: $73.1\% \pm 0.2\%$<br>Accuracy of Models 1 and 3: $69.4\% \pm 0.4\%$<br>Accuracy of Models 2 and 3: $76.8\% \pm 0.2\%$ | | |
| **Combined Error Rate** | Error Rate of Models 1 and 2: $11.0\% \pm 0.1\%$<br>Error Rate of Models 1 and 3: $9.4\% \pm 0.1\%$<br>Error Rate of Models 2 and 3: $11\% \pm 0.2\%$ | | |

**Figure 18:**
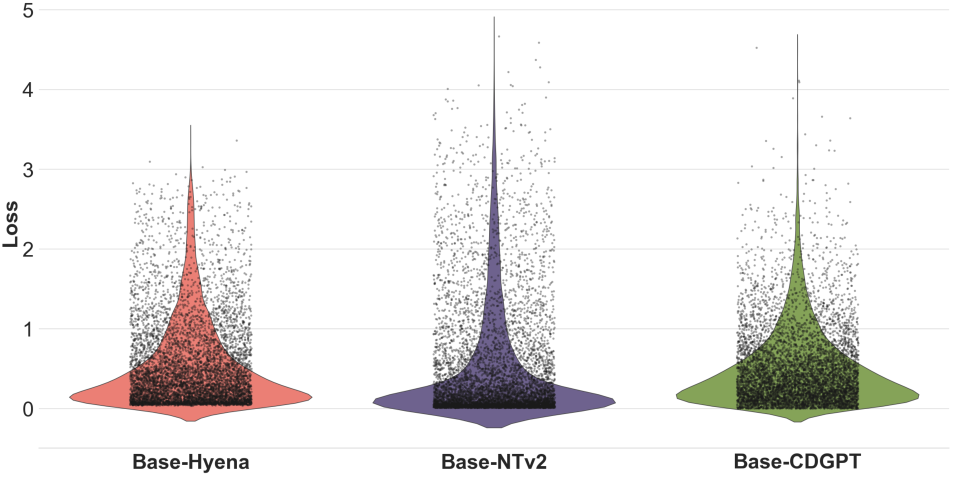
Short Sequence Experiment

**Figure 19:**
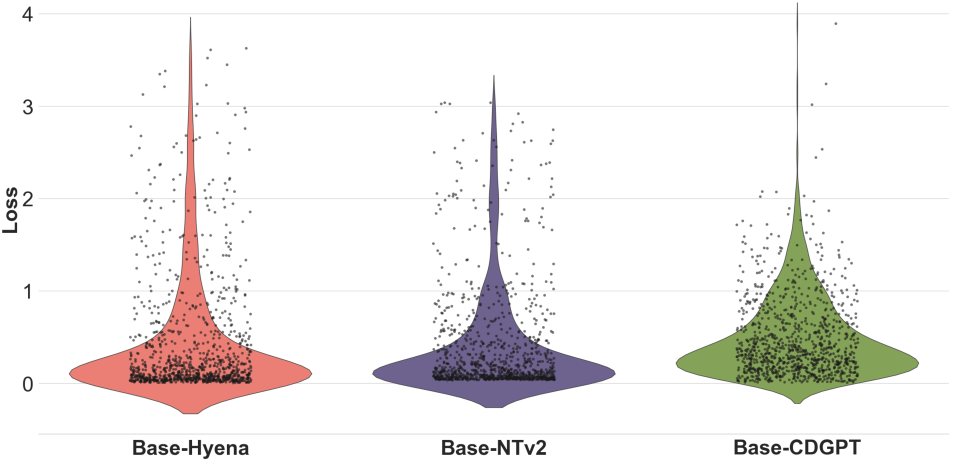
Long Sequence Experiment

**Figure 20:**
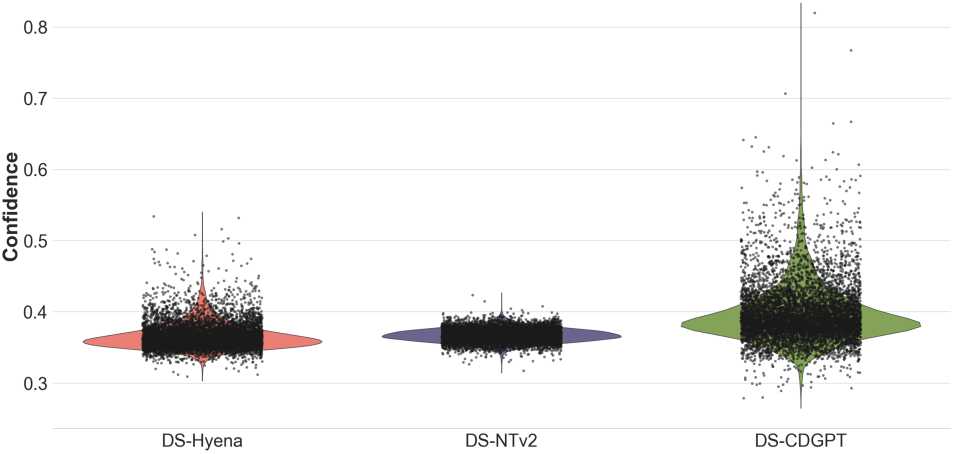
Short Sequence Experiment

**Figure 21:**
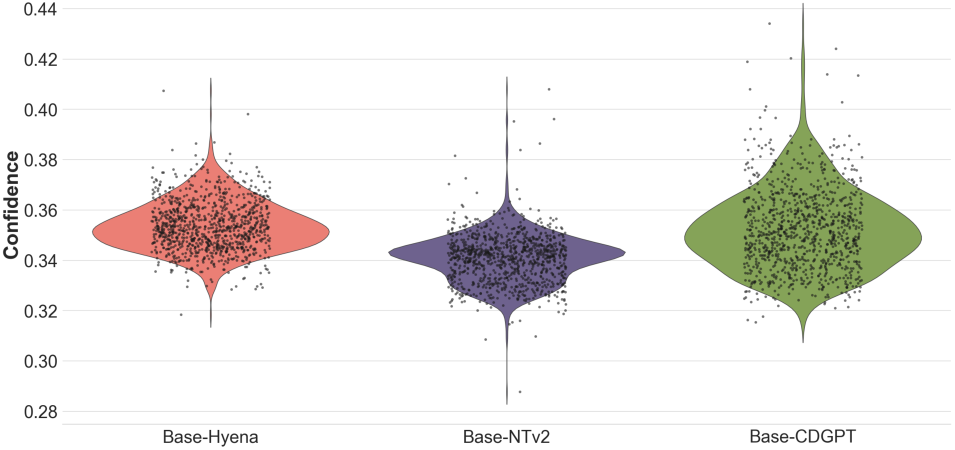
Long Sequence Experiment

